# Persistent Homology-based Functional Connectivity Explains Cognitive Ability: Life-span Study

**DOI:** 10.1101/2022.10.17.512619

**Authors:** Hyunnam Ryu, Christian G. Habeck, Yaakov Stern, Seonjoo Lee

## Abstract

Brain-segregation attributes in resting-state functional networks have been widely investigated to understand cognition and cognitive aging using various approaches (e.g., average connectivity within/between networks and brain system segregation). While these approaches have assumed that resting-state functional networks operate in a modular structure, a complementary perspective assumes that a core-periphery or rich club structure accounts for brain functions where the hubs are tightly interconnected to each other to allow for integrated processing. We introduce a novel method, persistent homology (PH)-based functional connectivity, to quantify the pattern of information during the integrated processing. We also investigate whether PH-based functional connectivity explains cognitive performance and compare the amount of variability in explaining cognitive performance for three sets of independent variables: (1) PH-based functional connectivity, (2) graph theory-based measures, and (3) brain system segregation. Resting-state functional connectivity data were extracted from 279 healthy participants, and cognitive ability scores were generated in four domains (fluid reasoning, episodic memory, vocabulary, and processing speed). The results first highlight the pattern of brain-information flow over whole brain regions (i.e., integrated processing) accounts for more variance of cognitive abilities than either brain system segregation or the graph theory-based network topology measure. The results also show that fluid reasoning and vocabulary performance significantly decrease as the strength of the additional information flow on functional connectivity with the shortest path increases.

## Introduction

he relationship between resting-state functional networks and cognition [11, 14, 22, 24], cognitive aging [50], and cognitive reserve [43] has been widely investigated. Brain functional connectivity has been quantified in various ways; for example, average correlation within/between networks [22, 24] and brain system segregation [11, 14]. These measurements are calculated based on the assumption that the brain network structure is modular. However, a complementary perspective assumes that a core-periphery or rich club structure accounts for brain functions where the hubs are tightly interconnected to each other to allow for integrated processing [2, 20, 38, 47].

Alternatively, graph theory-based analysis quantifies the topology of the brain network structure. For example, the segregation attribute can be measured by calculating cluster coefficient and local efficiency, or the integration attribute can be measured by calculating global efficiency and shortest path length [1, 29, 34]. However, these measures are calculated from a binary adjacency connectivity matrix obtained from thresholding at a specific value of the association measure. Thus, although the existing graph theory-based analysis helps researchers explore the effects of brain organization on cognitive functions and diseases, such results are difficult to reproduce and might critically depend on the thresholding choice [5, 39, 53].

Recent studies have used the minimum spanning tree (MST) method to address the thresholding dependency in graph theory-based analysis for the brain organization [5, 8, 25, 35, 45]. The MST method locates a unique spanning tree that connects N brain regions with N-1 edges at minimum cost (i.e., maximizing synchronization between brain regions). Under the assumption that the brain network is structured as a kind of transportation network, MST will serve as an important backbone of information flow in a weighted brain network [4, 35, 45, 46]. The MST considers topological efficiency (MST has the N-1 shortest path) and strength efficiency (MST has the highest connectivity strength among the possible trees). That is, MST constructs the backbone structure containing the strongest connections from the set of all available weighted connections that connect whole brain regions with the smallest number of edges.

Persistent homology (PH), one of the tools frequently used in topological data analysis (TDA) to study the shape of data, has been considered another way to address the thresholding issue of the brain network analysis [10, 13, 27]. The basic idea of TDA is to study data through their low-dimension topological features, which translate into connected components (dimension 0), loops (dimension 1), voids (dimension 2), and so on. To capture the low-dimension topological features, TDA builds an abstract structure upon the individual nodes in the network, which is called a simplicial complex. In a network structure, the simplicial complex consists of nodes and edges, like a subgraph (For the overall concept of TDA and a detailed explanation of simplicial complex with point cloud data, which is a basic data form in TDA, refer to [12, 16, 21]). There are many ways to construct the simplicial complex. A common method in network settings is to add edges to the nodes one by one in a way that the edge having a stronger weight is added first. Nodes are connected to each other with edges making connected components (dimension 1) or loops (dimension 2). If the network is even more complex, it is hard to find an appropriate number of edges added in the simplicial complex that will capture all of the topological features in the network. If, instead, all possible thresholding values are considered, the set of corresponding structures can capture the relevant topological features of the network. This is the focus of PH. The set of simplicial complexes defined as the edges added into the space is called filtration. PH tracks topological features that appear (birth) or disappear (death) as the filtration increases. In this way, PH captures topological features in the brain network, as well as avoids the thresholding issue. In fact, the algorithm for finding death filtration values (e.g., 1-correlation coefficients between node pairs) of dimension 0 PH (PH capturing connected components) in the network settings is the same as that for finding heights of the MST [26, 27]. That is, the MST is a tree composed of only edges having death filtration of dimension 0 PH, and dimension 0 PH has distribution information of edge weights of the structure with the most efficient information flow. From these characteristics, PH quantifies the pattern of functional connectivity of the MST network topology structure, which ensures the most efficient integrated processing of brain regions.

While the MST method only keeps the acyclic graph with no cycles (i.e., MST keeps only connected components with no loop structure), the PH method separately tracks the information of both connected components (dimension 0) and loops (dimension 1). Thus, PH can consider additional information through loops to MST before all nodes are connected. In addition, recent studies of MST network analysis in brain networks have viewed the constructed MST as a binary network and calculated the topological characteristics of the MST, such as the number of leaves (nodes with degree 1), the maximum betweenness of nodes, etc. [5, 8, 35, 39, 45]. From this perspective, MST network analysis differs slightly from the PH approach. MST network analysis quantifies the way the nodes in MST are arranged, while PH, when dimension 0 is only considered in the network setting, quantifies the edge weight pattern assigned at MST and loops added to MST.

In the current study, we call the tree connecting all brain regions with no loops (i.e., MST) and additional loops of this structure “backbone” and “cycle,” respectively. We introduce measures describing the distribution of weights in the backbone structure and cycle structure and call these measures PH-based functional connectivity. Briefly, the backbone structure describes brain-integrated processing, and the cycle structure describes how the brain regions are interconnected during brain-integrated processing. Thus, the distribution of weights in the cycle structure when the backbone structure is given can explain how tightly the hubs are interconnected to each other during brain-integrated processing. We also investigate the relationship between PH-based functional connectivity and cognitive aging. In addition, we compare the amount of variability in cognitive ability explained by PH-based functional connectivity to that explained by measures describing brain organization (network topology and MST), average functional connectivity, and brain system segregation.

## Persistent Homology based Functional Connectivity

### Multi-Scale Simplicial Complex

Rips filtration is the most natural way to construct a set of multi-scale simplicial complexes due to its computational efficiency [6]. Many TDA applications of brain networks have also used Rips complex filtration to calculate PH. Several studies have directly applied Rips complex filtration using the distance between nodes (for example, the distance between nodes can be defined as 1 – *r* (or 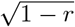), when the weighted network is constructed using Pearson correlations (*r*)) [18, 27, 28, 37, 44]. However, they considered only dimension 0 topological features (i.e., connected components). This is because the definition of dimension 1 topological features (i.e., loops) under Rips simplicial complex only covers non-clique cycles (for example, a triangle (3-clique) is a connected component, not a loop in Rips simplicial complex), which is not easily observable in a complex network. Here, to obtain a more interpretable and observable loop structure of a complex functional brain network, we use the graph simplicial complex, which is similar to the Rips simplicial complex but considers only 0- and 1-simplices (i.e., nodes and edges), excluding 2- and 3-simplices (i.e., filled triangles and tetrahedrons) and higher-order simplices from Rips complexes (Figure 1). Although graph filtration does not consider higher than order two relationships between nodes as connected components, we can use graph filtration to keep information of edges making all kinds of cycles (i.e., triangles, rectangles, pentagons, etc.), which describe the relationship between more than three nodes.

**Figure 1.**
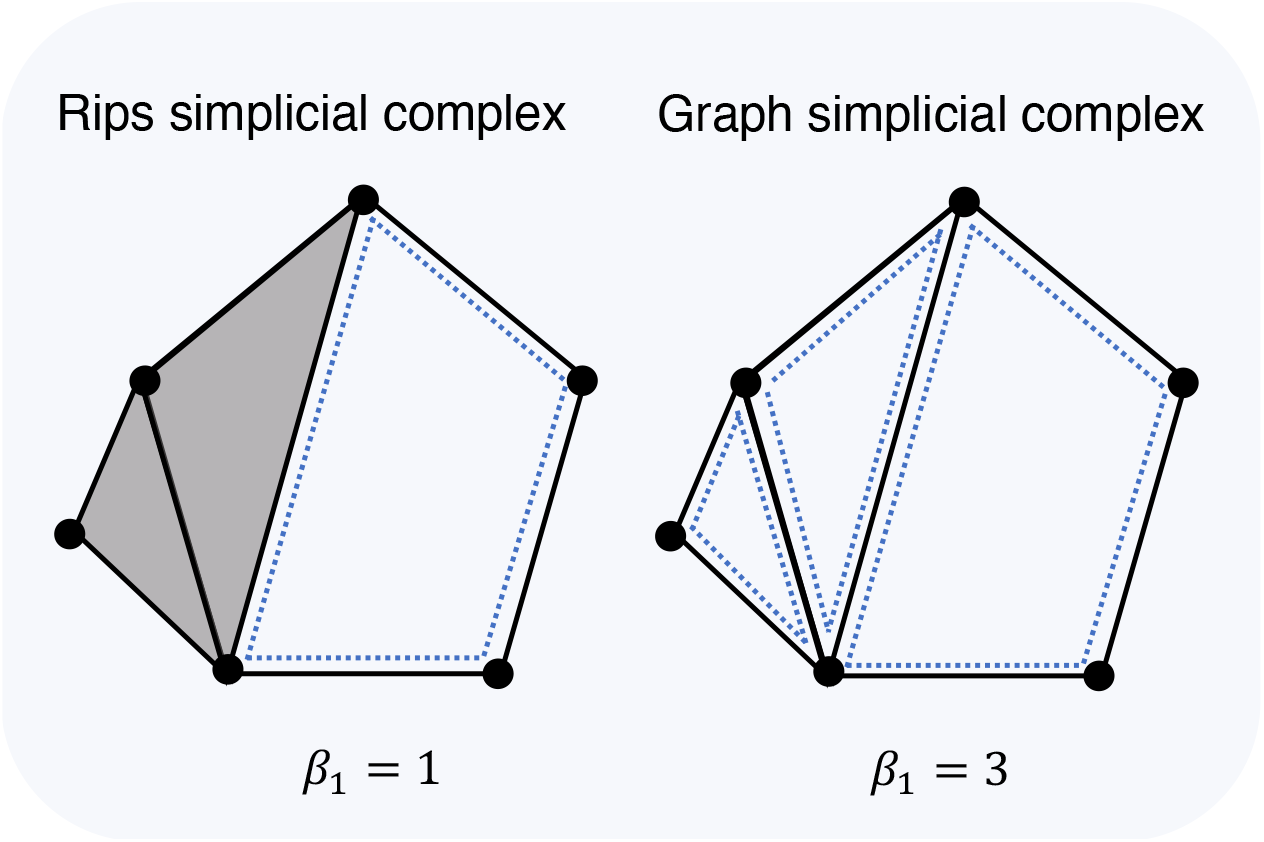
Example of Rips and graph simplicial complex. Rips simplicial complex (left) assigns grey faces to 3-cliques (triangles) and so has one loop (*β*_1_ = 1). Whereas graph simplicial complex (right) only considers points and edges and so has three loops in this example (*β*_1_ = 3)

### Algorithms for Graph Filtration and Persistent Homology

To construct the graph filtration, we used the distance between nodes as Rips filtration does. The algorithm for Rips filtration is to assign edges between two nodes of which distance is shorter than the filtration distance as the distance increases. In fact, this algorithm is the same as the minimum spanning tree (MST) algorithm as far as PH only considers connected components [26, 27]. MST can be created by sorting all edges in the network from the highest weight edge to the lowest weight edge (from the shortest distance edge to the longest distance edge), assigning one at a time, and keeping edges that make trees with no cycle [25]. To address cycles, the algorithm of the graph filtration keeps all edges in the MST algorithm, even if edges create cycles, but saves them separately (tree-making edges/cycle-making edges). Figure 2 illustrates how to construct the graph filtration with a toy example. With four nodes in the example, six edges (=4 × 3/2) are sorted on their weights in a decreasing way. When an edge is added one at a time, the edge assumes one of two roles: 1) it connects nodes that have not yet been connected so that one of the connected components dies (red solid edge in Figure 2) or 2) it connects already connected nodes, creating a new cycle (grey dashed edge in Figure 2). Before any edge is added, all nodes are connected components by themselves; thus, the number of connected components begins by the number of nodes (N) and becomes one connected component connected by N-1 edges. That is, each connected component has a birth time of zero and dies at the weight of that edge when each edge is added. Similarly, the added edge takes one of the two roles; therefore, out of the total N(N-1)/2 edges, except for the N-1 edge that reduces the connected component, the remaining (N-1)(N-2)/ 2 edges are involved in creating cycles. Here, each cycle has the birth time of the edge weight when the edge is included and does not die even when all edges are added. Therefore, each edge weight is included in either the death weight set of dimension 0 topological features or the birth weight set of dimension 1 topological features. In a formal definition,

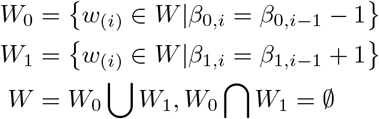

where *W*_0_ is a death weight set of connected components, *W*_1_ is a birth weight set of cycles, W is an edge weight set, *w*_(*i*)_ is the *i^th^* highest edge weight, and *β_k,i_* (*k* = 0,1) is the dimension *k* Betti number (i.e., the number of dimension k topological features) of the simplicial complex when edge with the weight *w*_(*i*)_ is added. The birth time set of dimension 1 topological features, *W*_1_, can also be decomposed as cycle weights before and after all nodes are connected, as below:

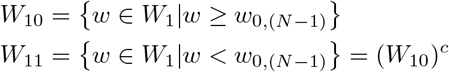

where *W*_10_ is a birth weight set of cycles born before all nodes are connected, *W*_11_ is a birth weight set of cycles born after all nodes are connected, and *w*_0,(*N*–1)_ is the minimum death edge weight of dimension 0 topological features.

**Figure 2.**
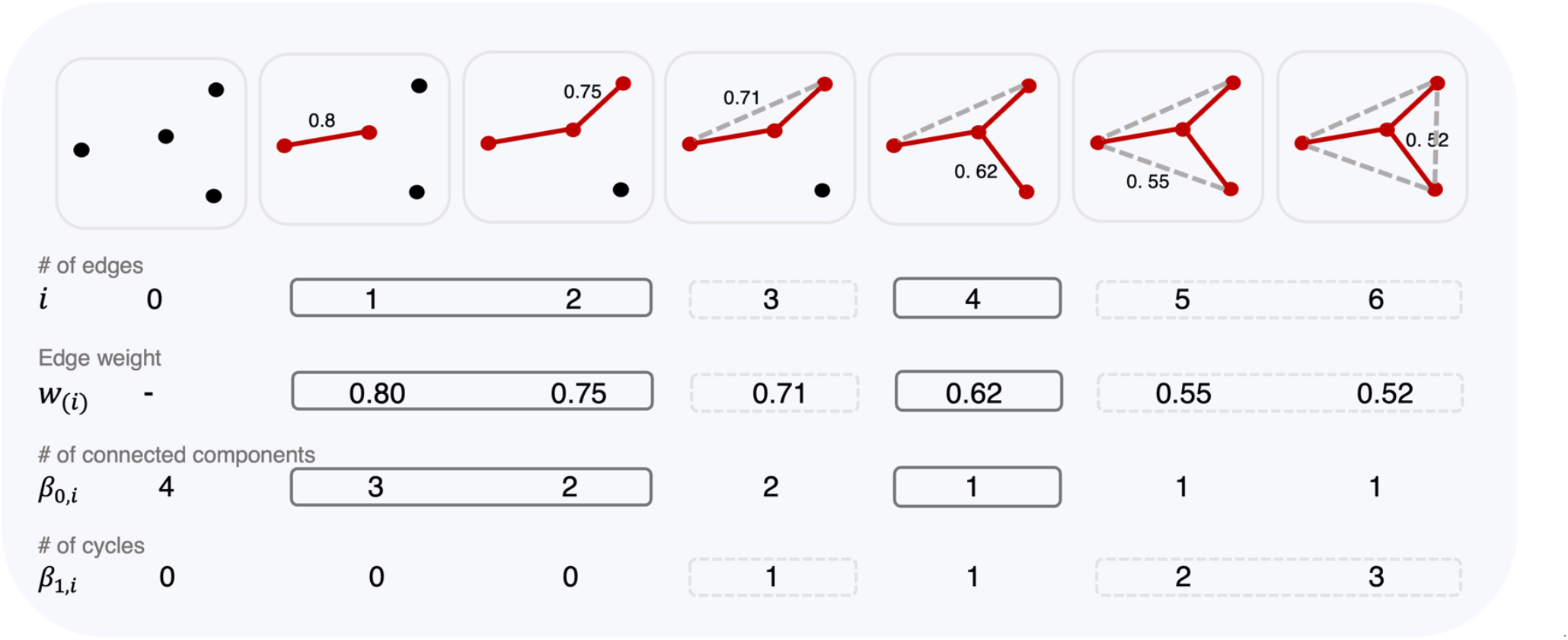
An example of graph filtration: Graph filtration consisting of four nodes and six edges. Each time a solid red edge is added, the number of connected components (*β*_0_) decreases by 1, and each time a dashed grey edge is added, the number of cycles (*β*_1_) increases by 1.

### PH-based Functional Connectivity

PH calculates the number of topological features across a whole range of filtration values (i.e., all possible edge weights in the network). Simultaneously, PH stores edge weight information in *W*_0_ (for dimension 0) and *W*_10_ (for dimension 1) when the number of topological features increases (for dimension 0) or decreases (for dimension 1) by one. While doing it, at some point, all nodes are connected into one connected component, which is the tree (i.e., no loops) connecting all N nodes with N-1 edges, of which weights are in *W*_0_ (i.e., MST). The studies related to MST have described the structure having MST as the “backbone” of the information flow [4, 35, 46], and here, we also called this structure “backbone.” Also, when edges with weights in *W*_10_ are assigned to the network, additional triangles are created upon the backbone structure, and we called these edges with weights in *W*_10_ “cycle.”

To describe the distribution of functional connectivity in the backbone and cycle structures with one value across a whole range of filtration values, we summarized the weight sets *W*_0_ and *W*_10_ with three measures: 1) backbone strength, defined as the average of elements in *W*_0_; 2) backbone dispersion, defined as the range (the largest value minus the smallest value) of elements in *W*_0_; and 3) cycle strength, defined as the average of values in *W*_10_. That is, backbone strength and cycle strength describe the mean functional connectivity of the brain backbone structure and the cycle structure, respectively. Since the backbone structure describes brain-integrated processing, and the cycle structure describes how the brain regions are interconnected during brain-integrated processing, interpretation of both backbone strength and cycle strength in the brain network can explain how tightly the hubs are interconnected to each other during brain-integrated processing. For example, as shown in Figure 3, there are cases where two resting-state brain networks have the same backbone strength but significantly different cycle strengths. Such information cannot be found only with dimension 0 PH (or MST), and additional cycle information in the backbone structure provides important information about the pattern of information flow. In Figure 3, A and B (C and D) visualize the functional network of an individual with the cycle strength at the first (third) quantile. At the same level of backbone strength, brain networks with high cycle strength have more cycles being involved in the flow of information when information begins to flow (i.e., with high functional connectivity, in this example, we only considered when the weight is higher than 0.6). Therefore, even with a similar backbone strength, low cycle strength is interpreted as information flow patterns with high strength are transmitted through the backbone (like highways) rather than additional cycles (like rural roads).

**Figure 3.**
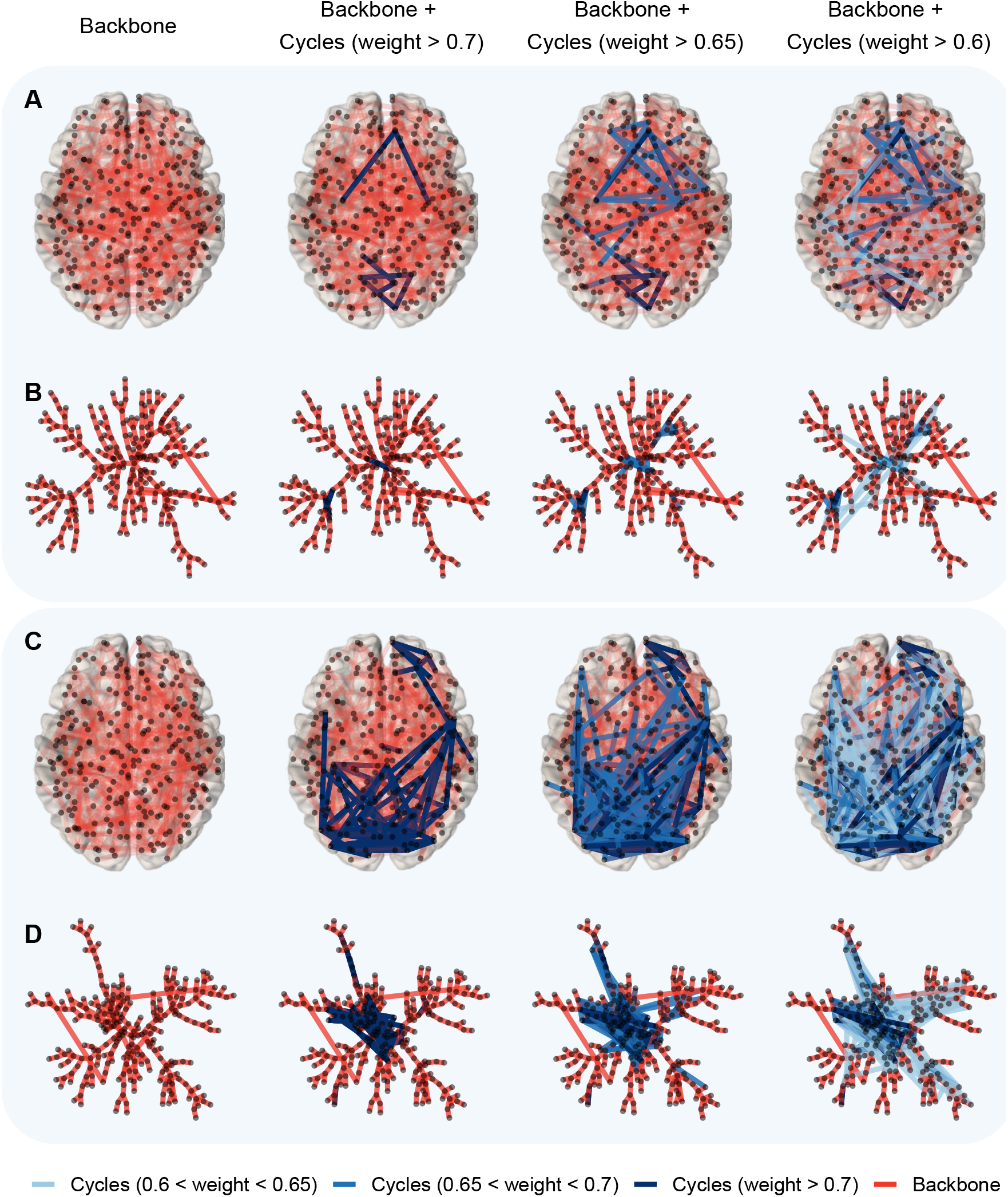
Visualization of functional connectivity illustrating backbone structure and additional cycle structure of two individuals. (**A**, **B**) Functional connectivity of individual who has 1st quantile cycle strength; (**C**, **D**) functional connectivity of individual who has 3rd quantile cycle strength. **A** and **C** show a top view of brain connectivity with 264 power ROIs’ MNI coordinates, and **B** and **D** visualize a tree-like graph layout based on a minimum spanning tree. The red line represents an edge in the backbone structure, and blue-tone lines represent edges making additional cycles (Darker blue represents a cycle with stronger edge weight). The first column visualizes only the backbone structure, and the second to fourth columns visualize the cycles in which weights are added in descending order. Only cycles having weights greater than the average backbone strength of 0.6 are shown in this Figure. Both individuals have the same backbone strength.

## Result

### Demographics

The characteristics of 279 participants are displayed in Table 1. The participants were aged from 20 to 80, and the younger (ages 20-39) age group had relatively more females and thicker cortexes than the middle-aged (ages 40-59) and older (ages 60-80) age groups. The four domain z-score for cognitive abilities and the three PH-functional connectivity measures are also presented separately for three age groups.

**Table 1.** Characteristics of Participants. Means (standard deviations in parentheses) or frequencies (percentage of sample in parentheses) are reported for each sample.

|  | Overall<br>(N=279) | Younger Adults<br>(ages 20-39)<br>(N=74) | Middle-Aged Adults<br>(ages 40-59)<br>(N=68) | Older Adults<br>(ages 60-80)<br>(N=137) |
| --- | --- | --- | --- | --- |
| Age | 53.573(16.835) | 29.419(5.397) | 51.279(5.563) | 67.759(5.065) |
| Education | 16.222(2.354) | 16.041(2.529) | 16.206(2.315) | 16.328(2.285) |
| Gender (Female) | 159(57%) | 51(68.9%) | 32(47.1%) | 76(55.5%) |
| Mean Thickness | 2.549(0.116) | 2.633(0.104) | 2.564(0.093) | 2.497(0.104) |
| Fluid Reasoning | 0.016(0.793) | 0.577(0.772) | 0.013(0.770) | -0.285(0.642) |
| Episodic Memory | 0.015(0.960) | 0.635(0.701) | 0.023(0.949) | -0.324(0.923) |
| Vocabulary | 0.068(0.901) | -0.380(0.956) | 0.041(0.901) | 0.324(0.770) |
| Processing Speed | 0.050(0.815) | 0.634(0.759) | 0.169(0.671) | -0.326(0.702) |
| Backbone Strength | 0.647(0.036) | 0.648(0.042) | 0.648(0.035) | 0.647(0.033) |
| Backbone Dispersion | 0.485(0.047) | 0.493(0.053) | 0.493(0.043) | 0.478(0.044) |
| Cycle Strength | 0.504(0.035) | 0.503(0.041) | 0.507(0.034) | 0.503(0.033) |

### Performance of PH-based Functional Connectivity

We compared the performance of PH-based functional connectivity measures in explaining cognitive performance with MST-based, functional connectivity-based, and network topology measures. We fitted each linear model with each z-score of cognitive ability as a dependent variable and one of the measures (three PH-based functional connectivity, two MST-based measures, two functional connectivity-based measures, and three network topology measures) as a predictor variable controlling for age, education, gender, and mean cortical thickness. Figure 4A displays the partial *R*^2^ of PH-based functional connectivity measures, MST-based measures, and connectivity-based measures with red-tone, blue-tone, and grey-tone bars, respectively. For fluid reasoning and vocabulary, cycle strength significantly explained variation of cognitive abilities (p <0.010) with higher partial *R*^2^ than all measures of MST and functional connectivity. For episodic memory, cycle strength also had the significant and highest partial *R*^2^ (p <0.010) among all measures, but backbone strength and overall functional connectivity also significantly explained the variation in cognitive ability (p <0.050).

**Figure 4.**
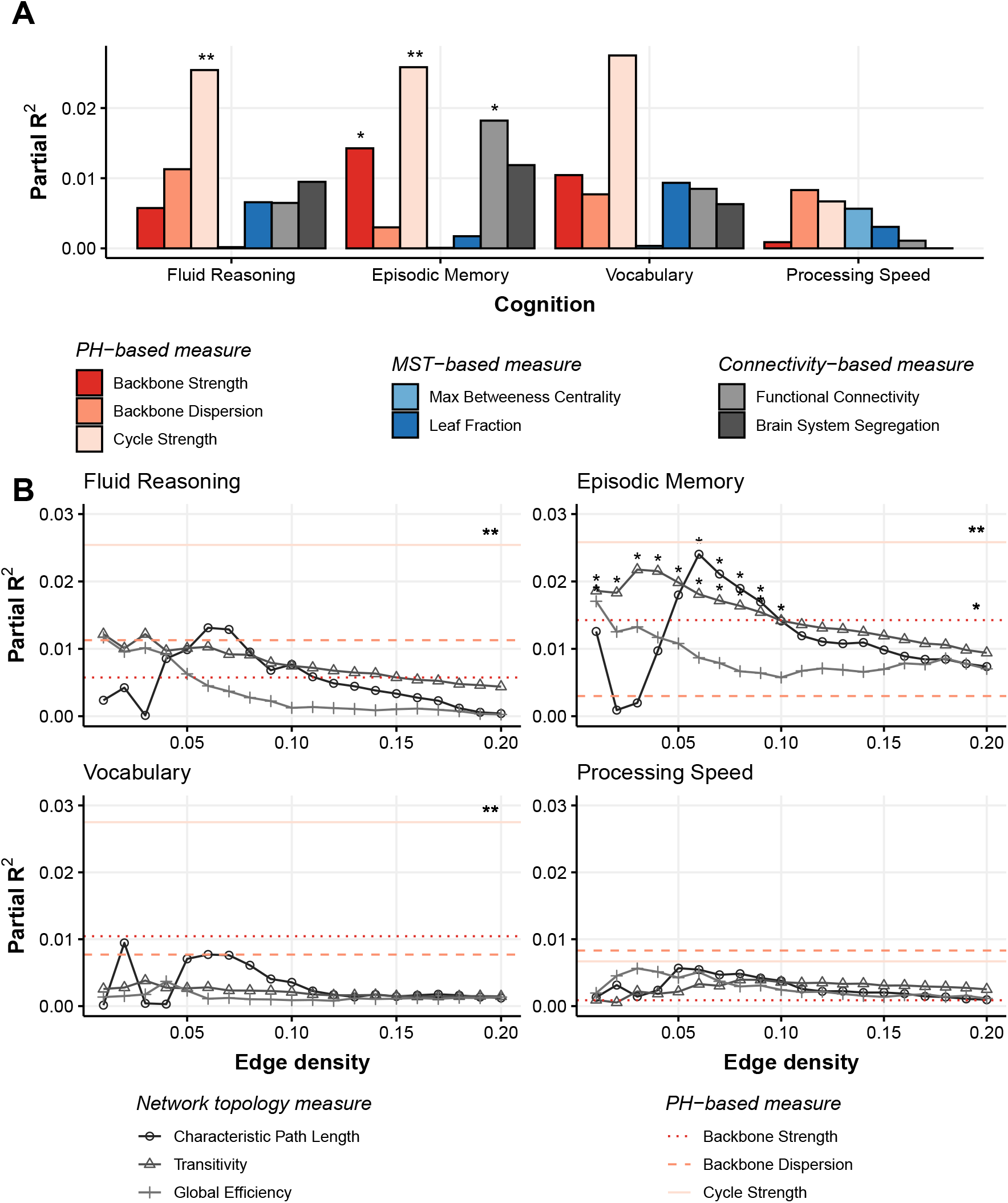
**A** Partial *R*^2^ comparison of PH-based measures with MST-based measures and connectivity-based measures in relation to four cognitive endpoints. The partial *R*^2^ is controlled by age, education, gender, and mean cortical thickness in each linear model. The partial *R*^2^ of PH-based functional connectivity measures, MST-based measures, and connectivity-based measures are represented with red-tone, blue-tone, and grey-tone bars, respectively. **B** Partial *R*^2^ comparison of PH-based measures with network topology measures at each thresholding value (0.01 to 0.20 by 0.01). The partial *R*^2^ is controlled by age, education, gender, and mean cortical thickness in each linear model. Grey-tone points represent partial *R*^2^ of network topology measures at each thresholding value. Red-tone horizontal lines represent partial *R*^2^ of PH-based functional connectivity measures across the whole range of thresholding values. (*p<0.050, **p<0.010, ***p<0.001)

Since graph theory-based network topology measures were different according to the thresholding values, we compared the performance of PH-based measures in explaining cognitive performance with graph network topology at each edge density. Figure 4B displays the partial *R*^2^ of PH-based functional connectivity measures and network topology measures calculated at each edge density as a thresholding value from 0.01 to 0.20 increased by 0.01. Grey-tone points represent partial *R*^2^ of network topology measures at each thresholding value. Red-tone horizontal lines represent partial *R*^2^ of PH-based functional connectivity measures across the whole range of thresholding values. Figure 4B shows that cycle strength significantly explains variation in the cognitive ability of fluid reasoning, vocabulary, and episodic memory (p<0.010), and backbone strength significantly explain variation in episodic memory cognitive ability (p <0.050), as shown in Figure 4A. Network topology measures at some point of thresholding value less than 0.1 (transitivity at 0.01 to 0.09, characteristic path length at 0.05 to 0.09, and global efficiency at 0.01) significantly explain variation in episodic memory (p<0.050). However, networks from our data set are not “complete” (all brain regions are not connected), with edge density less than 0.1, and in fact, Figure 4B shows that the results of partial *R*^2^ are not stable at thresholding less than 0.1. After thresholding 0.1, the results of partial *R*^2^ become stable but not significant.

### Effects of PH-based Functional Connectivity

Table 2 presents the correlation coefficients between PH-based functional connectivity and two demographic variables: age and cortical thickness. Among the three PH-based functional connectivity measures, the brain backbone dispersion was related to both age (*β*=-0.137, 95% CI −0.254 to −0.020, p<0.05) and mean cortical thickness (*β*=0.217, 95% CI 0.102 to 0.332, p<0.001); the relationship between brain backbone dispersion and mean cortical thickness was significant even after multiple comparisons correction. As a posthoc test, we also conducted a mediation analysis for brain backbone dispersion to examine a potential mediating role of cortical thickness in the relationship between age and backbone dispersion. Figure 5 displays the result of the mediation analysis. The mediation analysis was implemented with a bootstrapping process to construct confidence intervals for the indirect effect and proportion of mediated effect[32]. The mean cortical thickness fully mediated the effect of age on brain backbone dispersion (standardized indirect effect = −0.110, 95% CI −0.186 to −0.040, p<0.010), and the proportion of the mediated effect of mean cortical thickness was 0.802 with 95% CI 0.176 to 4.310 (p<0.050).

**Table 2.** Correlation between Age and Cortical Thickness, and PH-based functional connectivity measures

| Demographics | PH-measure | Standardized Correlation Coefficients | [95% CI] | P-value |
| --- | --- | --- | --- | --- |
| Age | Backbone Strength | -0.016 | [-0.134,0.102] | 0.791 |
| Age | Backbone Dispersion | -0.137 | [-0.254,-0.020]* | 0.022 |
| Age | Cycle Strength | -0.004 | [-0.122,0.113] | 0.941 |
| Cortical Thickness | Backbone Strength | -0.011 | [-0.128,0.107] | 0.860 |
| **Cortical Thickness** | **Backbone Dispersion** | **0.217** | **[0.102,0.332]\*\*\*** | **<0.001** |
| Cortical Thickness | Cycle Strength | -0.031 | [-0.149,0.087] | 0.607 |

\*p<0.050, \*\*p<0.010, \*\*\*p<0.001.
Statistically significant values after FDR correction are bolded.

As shown in Figure 4A, overall functional connectivity appears to account significantly for the variability of episodic memory performance, as well as backbone strength and cycle strength. This means that the effect of backbone strength and cycle strength may be affected by overall functional connectivity. Therefore, in order to investigate the association between cycle strength and cognitive abilities regardless of overall functional connectivity level, a regression model was fitted, controlling the overall functional connectivity along with age, gender, education, and mean cortical thickness. Table 3 presents the linear regression results using each z-score in four domains of cognitive performance as a dependent variable and one of three PH-based functional connectivity measures as a predictor after controlling age, education, mean cortical thickness, and overall functional connectivity. Cycle strength was negatively associated with fluid reasoning (*β*=-0.198, 95% CI −0.348 to −0.047, p<0.050), and vocabulary (*β*=-0.204, 95% CI −0.363 to −0.044, p<0.050) and these relationships survived after multiple comparisons correction.

**Table 3.** Linear regression between cognitive performances and PH-measures

| Cognitive Performance | PH-measure | Standardized Correlation Coefficients | [95% CI] | P-value |
| --- | --- | --- | --- | --- |
| Fluid Reasoning | Backbone Strength | -0.002 | [-0.282, 0.279] | 0.991 |
| Fluid Reasoning | Backbone Dispersion | 0.092 | [-0.007, 0.191] | 0.071 |
| **Fluid Reasoning** | **Cycle Strength** | **-0.198** | **[-0.348, -0.047]\*** | **0.011** |
| Episodic Memory | Backbone Strength | 0.054 | [-0.244, 0.351] | 0.724 |
| Episodic Memory | Backbone Dispersion | 0.053 | [-0.053, 0.16] | 0.325 |
| Episodic Memory | Cycle Strength | -0.122 | [-0.284, 0.039] | 0.139 |
| Vocabulary | Backbone Strength | -0.114 | [-0.41, 0.182] | 0.450 |
| Vocabulary | Backbone Dispersion | 0.081 | [-0.024, 0.187] | 0.132 |
| **Vocabulary** | **Cycle Strength** | **-0.204** | **[-0.363, -0.044]\*** | **0.013** |
| Processing Speed | Backbone Strength | 0.010 | [-0.278, 0.298] | 0.945 |
| Processing Speed | Backbone Dispersion | 0.080 | [-0.022, 0.182] | 0.127 |
| Processing Speed | Cycle Strength | -0.117 | [-0.273, 0.039] | 0.142 |

All regressions controlled for age, gender, education, mean cortical thickness, and overall functional connectivity.
+ $p < 0.100$ , \* $p < 0.050$ , \*\* $p < 0.010$ , \*\*\* $p < 0.001$ .
Statistically significant values after FDR correction are bolded.

## Discussion

### PH-based Functional Connectivity Explains Cognitive Abilities

In neuroscience, researchers have used various methods to describe how the brain network functionally works during the performance of cognitive tasks or rest. For example, recent studies investigating the relationship between brain functional connectivity and cognitive abilities have mainly utilized average functional connectivity (over the whole brain or within or between networks) [14, 22, 24], the brain system segregation [11], and graph theory-based network topology measures [29]. In the current study, we introduced persistent homology (PH)-based functional connectivity measures and applied these measures to investigate how functional connectivity in the most efficient information flow structure explains cognitive abilities. PH-based functional connectivity accounts for the pattern of information flow when all brain regions are connected into one connected component (i.e., integration) in the most efficient way.

**Figure 2.**
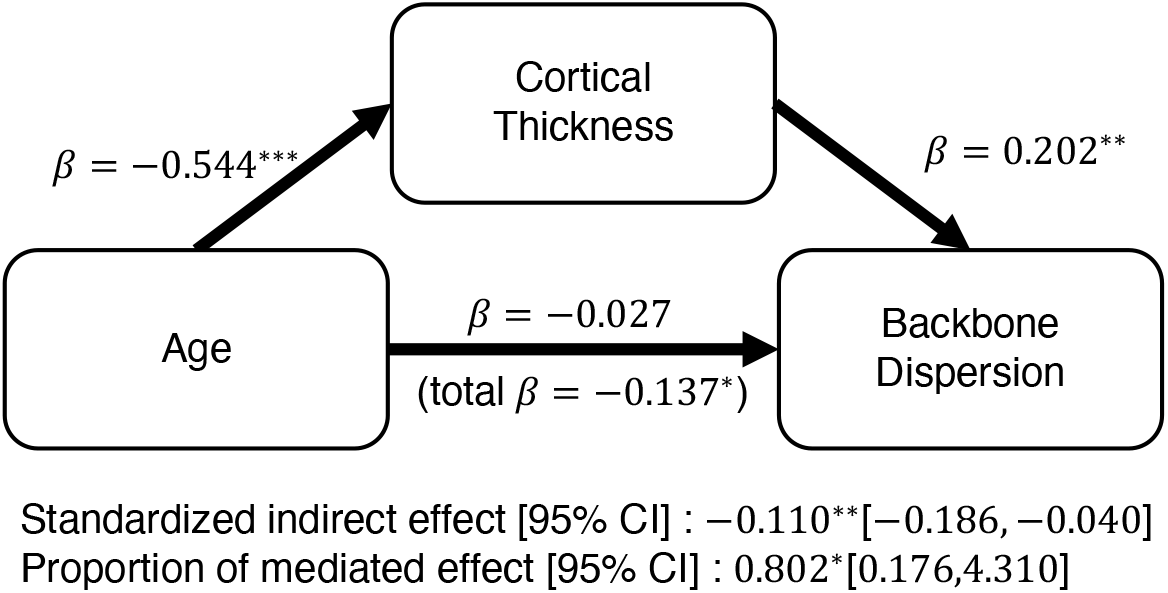
Mediation analysis. The path diagram (including standardized coefficients and significances) of the mediation analysis demonstrates that cortical thickness fully mediates the effect of age on backbone dispersion. (*p<0.05, **p<0.01, ***p<0.001)

From that point of view, there are several advantages of PH-based functional connectivity compared to the existing brain functional connectivity measures. Firstly, PH-based functional connectivity is robust to weak, noisy connectivity. Overall functional connectivity, which is calculated by averaging all connectivity values across the whole brain, tends to be biased downwards due to many weak connectivity values. On the other hand, PH-based functional connectivity only considers the edges involved in connecting all brain regions efficiently and ignores the edges with connectivity after all brain regions are connected. Thus, PH-based functional connectivity can relieve the effect of weak, noisy edge weights. Secondly, PH-based functional connectivity describes brain-integrated processing of information flow. The existing methods describing brain functional connectivity, such as functional connectivity within/between networks or brain system segregation, assume that brain network structure is modular. However, it has often been hinted that the brain integration attribute may play as important a role in explaining brain behavior as the segregation attribute [20, 47]. In previous studies, overall functional connectivity was mainly used as a measure of brain integration properties. But, as mentioned earlier, overall functional connectivity is highly likely to be underestimated in describing brain integration attributes. On the other hand, PH-based functional connectivity measures can explain the integration properties more intuitively because they consider the tree connecting all brain regions. Lastly, persistent homology is free from the thresholding issue that has been frequently pointed out in the graph theory-based network analysis [5, 39, 49, 53]. Thus, PH-based functional connectivity has an advantage in that it can bring consistent analysis results regardless of the thresholding choice [10, 13, 27]. We also found that results from topology network measures based on graph theory were unstable when the network was quite sparse, while PH-based functional connectivity measures are the same across the whole range of filtration (see Figure 4B).

When looking at the performance of PH-functional connectivity in explaining cognitive abilities, the results in Figure 4 show that PH-based functional connectivity performed better in explaining cognitive ability than overall functional connectivity, brain segregation, or graph theory-based measures. These results suggest that cognitive abilities have more to do with the pattern of information flow over the whole brain than with connections within and between the networks.

### Edge Weight Distribution of Brain Backbone and Cortical Thickness

One of the main concepts of PH-based functional connectivity in the current study is “backbone.” The algorithm for finding edges involving reducing the number of connected components (dimension 0 topological features) in the graph filtration PH setting is the same as MST algorithm. Since the union of all shortest paths coincides with the MST, the MST might represent the critical backbone of the information flow [8, 35, 48], and thus the death weight set of connected components, *W*_0_, has information of backbone weights. Among our PH-functional connectivity measures, backbone strength and backbone dispersion describe the distribution of weights of backbone structure. Backbone dispersion, which is the range of the distribution of *W*_0_, represents the strength difference in the critical backbone of the information flow. In the current study, backbone dispersion was positively associated with the mean cortical thickness (see Table 2 and Figure 5). This suggests that the greater the thickness of the grey matter cortex of the brain, the greater the difference in the strength of connections in the flow of information to the whole brain.

### Cycle Strength and Cognitive Abilities

Another main concept of PH-based functional connectivity in the current study is “cycles.” The cycle provides additional information to the backbone of information flow, while MST only considers a tree with no loops. In other words, the backbone strength represents the overall strength of the information flow, but the pattern of strength omitted in the process can be reinforced from the birth weight set of cycles, *W*_1_0. As shown in the example of Figure 3, even with a similar backbone strength, low cycle strength is interpreted as information flow patterns with high strength are transmitted through the backbone rather than additional cycles. In the current study, we found that fluid reasoning and vocabulary were negatively correlated to cycle strength (Table 3). This suggests that cognitive ability increases as the flow of important information (here, strong connectivity) tend to go through the backbone structure without being dispersed into several cycles.

### PH-based Functional Connectivity vs Betti Curve Features

Unlike general persistent homology on point cloud data, which is interested in how many significant hole structures appear from data points sampled from the underlying shape of the data, the topological features appearing in the brain connectivity describe a way of connecting predefined nodes (i.e., brain regions). Thus, to effectively describe these characteristics of topological features of brain connectivity, the Betti curve, which tracks the number of topological features (e.g., connected components or loops) at each filtration value, has been considered one of the summarized representations of the persistent homology of brain connectivity. Many applications of the Betti curve to brain connectivity have used measures describing the shape of the Betti curve such as the area under the Betti curve and the slope of the Betti curve [18, 26, 27, 44]. These summaries of Betti curves quantify the shape characteristics of Betti curves rather than directly summarizing the dynamic topological characteristics of brain connection structures. For example, while the slope of the dimension 0 Betti curve has been interpreted as the “speed” of brain connectivity, the concept of “speed” here is very abstract and derivative. “Speed” here mathematically means the change in the number of connected components as the filtration value (i.e., Pearson correlation) changes. In fact, the topology referred to in graph theory-based analysis focuses on how nodes are arranged or interconnected, whereas the topology referred to in persistent homology (hence, Betti curves) refers to the number of holes in the space. In addition, unlike graph theory-based analysis that quantifies the network topology varying with filtration (or thresholding), the Betti curve is constructed in a way that captures the filtration values where birth and death occur in each Betti number. Therefore, the interpretation of the Betti curve is more natural when focusing on the filtration values according to each fixed Betti number.

In the current study, we applied persistent homology with graph filtration to decompose the weight set into dimension 0 death weight set and dimension 1 birth weight set. In fact, the application of graph filtration can make the individual Betti curve only depend on the edge weights when the connected component dies and the cycle is born. Thus, instead of using the Betti curve, we considered the weight sets of edges involving the death of each connected component and the birth of each cycle. The weights included in *W*_0_ are the same as the death filtration observed for each dimension 0 Betti number. By doing so, we sought to extend the interpretation of PH from simply describing the speed of brain connectivity to explaining the cost distribution of the economic backbone structure (i.e., MST) and cycles adding information to MST of the whole brain.

Moreover, using the defined weight set, the dimension 0 Betti curve can be defined as follows:

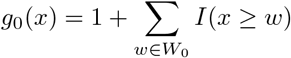

where *I*(·) is an indicator function, and *g*_0_(*x*) is dimension 0 Betti curve. Also, the standardized Betti curves are defined as:

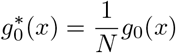

where *N* is the number of nodes, 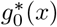 is the standardized dimension 0 Betti curve. Then, the area under the standardized Betti curve is negatively related to the mean of 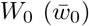, which is backbone strength in this article, as follow:

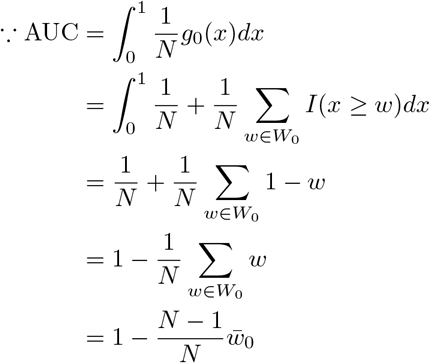

Therefore, the small value of the area under the curve (AUC) of dimension 0 Betti curve can be interpreted as the high average strength of the backbone structure (i.e., the overall strength of information flow in the brain network is high).

As in recent studies related to the application of PH to the brain network, the summary values (e.g., AUC) of the Betti curve considering only dimension 0 consider limited edge weight information related to the backbone, so it may not be sufficient to explain the pattern of brain functional connectivity in some cases. At least in this study, the overall strength of brain information flow, for example, the backbone strength (or AUC of dimension 0 Betti curve), was insufficient to explain cognitive ability. Instead, additional information on cycle structure in the current study suggested that cognitive ability is related to patterns of brain information flow.

## Conclusions

In sum, in this study, we presented a novel measure to quantify an important aspect of functional connectivity which characterizes the pattern of whole-brain integration by using PH. We applied the new measures to study the relationship between the pattern of resting-state brain functional connectivity and cognitive ability. Our approach offers a novel insight into brain functional connectivity during integrative processing, and PH-based functional connectivity measures explain cognitive aging, while segregation-focused connectivity measures or graph theory-based network topology does not.

## Materials and Methods

### Resting-state fMRI Data Acquisition

The resting-state fMRI data used in this article were obtained from Reference Ability Neural Network (RANN) Study and Cognitive Reservation (CR) Study at Columbia University Irving Medical Center [40, 41, 42]. The data set is composed of 279 participants (gender: 159 females, 120 males; age: 53.57 ± 16 (range 20-80)). A detailed description of the data set is available in previous reports [40, 42, 51].

MRI data were collected on a 3.0T Philips Achieva Magnet. There were two, 2-hr MR imaging sessions to accommodate the twelve fMRI activation tasks as well as the additional imaging modalities. At each session, T1-weighted MPRAGE scan (repetition time (TR) = 6.5 ms, echo time (TE) = 3 ms, flip angle = 8°, field of view = 254 mm × 254 mm, in-plane resolution = 256 × 256, 165 - 185 axial slices, slice-thickness = 1 mm, gap = 0 mm) was obtained for later use in preprocessing. Resting-state fMRI blood oxygen level-dependent (BOLD) scans (TR = 2000 ms, TE = 20 ms, flip angle = 72°, in-plane resolution = 112 × 112, 37 axial slices, slice-thickness = 3 mm, gap = 0 mm, scan time = 9.5 min, 285 volumes) were conducted. The participants were instructed to lie still with their eyes closed and not think of anything in particular, while staying awake.

### Resting-state fMRI Data Preprocessing

Images were preprocessed using an in-house developed native space method [33]. Briefly, the preprocessing pipeline included slice-timing correction and motion correction (MCFLIRT) performed using the FSL package [23]. All volumes were registered (6 df, 256 bins mutual information, and sinc interpolation) to the middle volume. Frame-wise displacement (FWD) [30] was calculated from the six motion parameters and root-mean-square difference (RMSD) of the BOLD percentage signal in the consecutive volumes. To be conservative, the RMSD threshold was lowered to 0.3% from the suggested 0.5%. Contaminated volumes were then detected by the criteria FWD ¿ 0.5 mm or RMSD ¿0.3% and replaced with new volumes generated by linear interpolation of adjacent volumes. Volume replacement was performed before temporal filtering [9]. Volume replacement was performed before temporal filtering [15]. Flsmaths–bptf was used to pass motion-corrected signals through a bandpass filter with cut-off frequencies of 0.01 and 0.09 Hz. Finally, the processed data were residualized by regressing out the FWD, RMSD, left and right hemisphere white matter, and lateral ventricular signals [3]. Using advanced normalization tools (ANTs), each T1 image was registered to the 2 mm MNI template, and the residualized images were warped to the 2 mm MNI template.

### Cognitive Abilities

Twelve measures were selected from a battery of neuropsychological tests to assess cognitive functioning [36]. Fluid reasoning was assessed with scores on three different tests: Wechsler Adult Intelligence Scale (WAIS) III Block design task, WAIS-III Letter–Number Sequencing test, and WAIS-III Matrix Reasoning test. For processing speed, the Digit Symbol subtest from the WAIS-Revised [52]. Part A of the Trail making test and the Color naming component of the Stroop [17] test were chosen. Three episodic memory measures were based on sub-scores of the Selective Reminding Task [7]: the long-term storage sub-score, continuous long-term retrieval, and the number of words recalled on the last trial. Vocabulary was assessed with scores on the vocabulary subtest from the WAIS-III, the Wechsler Test of Adult Reading, and the American National Adult Reading Test [19]. Domain scores were generated by z-scoring performance on each task relative to the full study sample, then average z-scores were computed for tasks within each domain (four domain z-scores: Fluid Reasoning, Processing Speed, Episodic Memory, Vocabulary).

### Functional Connectivity

The nodes were defined using an atlas derived from [31] and the mean time series of each node was extracted. Due to the lack of whole cerebellum coverage during resting-state scans, only non-cerebellar ROIs were included in our analyses (264 ROIs - 8 cerebellar ROIs = 256 ROIs in total). Pearson correlations were then performed for all pairwise combinations of ROIs. Thus, 32,640 (= 256 × 255 / 2) functional connectivity pairs (or edges) were used in our analysis. In this article, we use absolute Pearson correlations to take both positive and negative synchronization of brain signals into account.

### Global network topology measures

Since PH-based measures quantify the strength of whole-brain (i.e., global) backbone organization, we considered three selected global network topology measures to be compared with PH-based measures [34]: 1) characteristic path length, defined as the average shortest path length between all nodes; 2) transitivity, defined as the relative number of triangles in the graph, compared to the total number of connected triples of nodes; and 3) global efficiency, defined as the average reciprocal of the shortest path length between all nodes. The characteristic path length and global efficiency are related to network organization efficiency, and transitivity is related to the segregation of network organization. Since network topology measures differ according to the thresholding selection, we calculated the network topology measures at each edge density (from 1% to 20%).

### MST-based measures

We used two global MST-based measures [35]: 1) maximum betweenness, defined as the maximum number of shortest paths through each node in a graph and 2) leaf fraction, defined as the fraction of the number of nodes with degree 1 among the number of all possible edges. We can quantify the configuration of MST with maximum betweenness and leaf fraction. High maximum betweenness and high leaf fraction indicate that MST is more likely to appear as a star-type, while low maximum betweenness and low leaf fraction indicate line-type MST [45].

### Connectivity-based measures

We quantified individual resting-state functional connectivity with the mean Fisher z-transformed correlation coefficient of all pairs of nodes. In addition, we considered measures representing segregation of brain organization based on brain functional connectivity. We used the brain system segregation (BSS) [11], defined as follows:

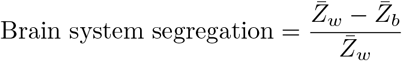

where 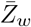 is the mean Fisher z-transformed correlation coefficient between nodes within the same system (defined by 31) and 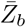 is the mean Fisher z-transformed correlation coefficient between nodes of one system to all nodes in other systems.

## Data, Materials, and Software Availability

R-package “PHfconn” (available at https://github.com/hyunnamryu/PHfconn) contains source code files to compute the persistent homology-based functional connectivity. The package also contains the example data and code files for statistical evaluations and visualizations presented in the article. Data are available upon reasonable request, and are subject to a formal data use agreement.

## Acknowledgements

This work was supported by three grants from the National Institute on Aging (R01AG026158, principal investigator: Y.S.; R01AG038465, principal investigator: Y.S.; R01AG062578, principal investigator: S.L.)

